# An innovative non-invasive technique for subcutaneous tumour measurements

**DOI:** 10.1101/622605

**Authors:** Juan Delgado-SanMartin, Beate Ehrhardt, Marcin Paczkowski, Sean Hackett, Andrew Smith, Wajahat Waraich, James Klatzow, Adeala Zabair, Anna Chabokdast, Leonardo Rubio-Navarro, Amar Rahi, Zena Wilson

## Abstract

In oncological drug development, animal studies continue to play a central role in which the volume of subcutaneous tumours is monitored to assess the efficacy of new drugs. Tumour volume is currently estimated by measuring length and width with callipers and then estimating the volume of the tumour as if it were a regular spheroid. However, this method is subjective, insufficiently traceable, and is subject to error in the accuracy of volume estimates as tumours frequently are irregular.

This paper explores the extent of inconsistencies in calliper measurements by conducting a statistical review of a large dataset consisting of 2,500 tumour volume measurements from 1,600 mice by multiple operators across 6 mouse strains and 20 tumour models. We also explore the impact of six different tumour morphologies on volume estimation and the detection of treatment effects using a computational tumour growth model. Finally, we propose an alternative method to callipers for estimating volume – BioVolume™, a 3D scanning technique. BioVolume simultaneously captures both stereo RGB (Red, Green and Blue) images from different light sources and infrared thermal images of the tumour. It detects the tumour region automatically and estimates the tumour volume in under a second. BioVolume has been tested on a dataset of 297 scans from over 120 mice collected by four different operators.

This work demonstrates that it is possible to record tumour measurements in a rapid, minimally invasive, morphology-independent way, and with less human-bias compared to callipers, whilst also improving data traceability. Furthermore, the images collected by BioVolume may be useful, for example, as a source of biomarkers for animal welfare and secondary drug toxicity / efficacy.

## 1 Introduction

Animal models of human cancers are fundamental to our understanding of tumour biology. Tumour volume is a significant metric for preclinical trials where it provides a surrogate measure of both disease progression and treatment efficacy. Thus, accurate and repeatable estimation of tumour volume is crucial to declare a given trial to be a success or failure with confidence (1). At present, it is standard practice to estimate subcutaneous tumour volume by using callipers to take manual measurements of tumour length and width. This approach erroneously assumes tumours to be regular spheroids (1,2), when they are proven to be irregular. Medical imaging technologies such as magnetic resonance imaging (MRI), computed tomography (CT) and ultrasound (US) offer an alternative but require the immobilisation of the animals (by anaesthesia) and are resource intensive, thereby potentially compromising animal welfare, increasing costs, and creating logistical complications (3–6). Efforts have been made to produce more accessible alternatives to these methods, such as 3D stereo photometry, time-of-flight and structured light (7–11).

In what follows, we explore how manual calliper measurements introduce human bias into pre-clinical trials. Furthermore, we use a cellular automaton model to investigate how calliper measurements influence our understanding of study outcomes for a range of different tumour morphologies (12). We propose an alternative measurement method, named *BioVolume™ (Fuel3D, Oxford, UK; www.fuel3d.com*) which combines 3D stereo photometry (to capture depth) and infrared/thermal imaging (to delineate the poorly vascularised tumour region).

## 2 Methods

### 2.1 Datasets

We analysed two datasets:

– <u>Dataset 1, Calliper statistical review:</u> Records for 1,608 mice and 2,488 calliper measurements were collected by 29 AstraZeneca operators over a period of 17 months from February 2017 until June 2018. The measurements belong to 43 pre-clinical studies.
– <u>Dataset 2, BioVolume – calliper comparison:</u> We collected scans of tumours from 120 mice using BioVolume on 3 occasions between 14/09/18 and 05/10/18. Calliper measurements were also taken for each tumour. A total of 257 calliper measurements and 297 scans were collected by four operators.

Five different strains of mice were sourced from Charles River UK (www.criver.com) & Envigo UK (www.envigo.com): SCID, BALB/c, C57/BL6, NSG and Nude. More information regarding the data can be found in Appendix 1 (S2.1).

### 2.2 BioVolume™

BioVolume is a small desktop device (27 x 18.5 x 16.8 cm), which captures both thermal (infrared) and 3D surface images. To acquire a scan, the mouse is held such that the tumour region is exposed to the device opening (see Figure 1, left). Then, acquisition is triggered on either a connected laptop or by using a button on the device itself (see Figure 1). There is no requirement to anesthetise the animal. Acquisition takes 0.25s, and rendering occurs in the cloud. Rendering a full image requires approximately 25s. This can be parallelised while the rest of images are acquired. The segmentation and measurement processing modules run in under a second, and measurements are displayed to the operator immediately.

**Figure 1:**
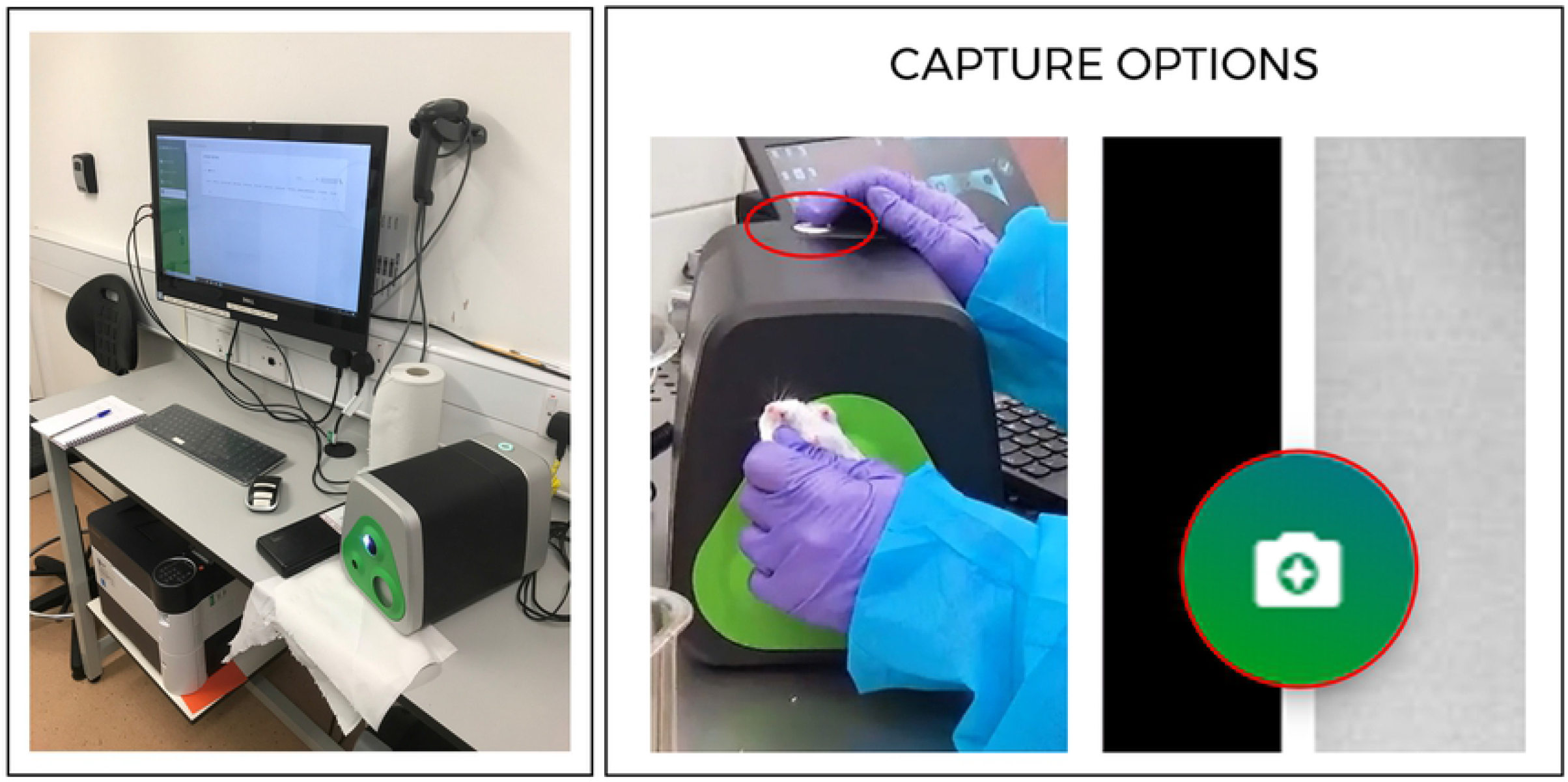
complete set up of the BioVolume unit including computer monitor and desktop device (left), closeup image of a white SCID mouse being presented to the aperture of BioVolume (right). Two capture options are shown.

A BioVolume unit consists a stereo system with two RGB cameras, three white light flashes and an infrared thermal camera. Upon activation, the unit collects 6 photographic (RGB) images and a thermal frame. The BioVolume software utilised in this work was a beta version named v0.1_f5bf15. The RGB images are reconstructed in a manifold by means of a binocular stereo-process, outputting both depth and RGB maps (13,14). These maps are then co-registered onto the thermal frame using a conventional transformation based upon a prior positional calibration of the RGB and thermal cameras, respectively. The segmentation of the tumour happens on the thermal map, which is then projected onto the depth map. The height is then obtained by fitting a plane to the back of the mouse using an optimisation algorithm. More details in Appendix 1.

### 2.3 Volume calculation

We compare two formulae for the estimation of tumour volume:

– Spheroid formula (BioVolume & callipers):

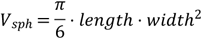
– Cylindrical volume (BioVolume):

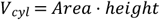

### 2.4 Computational Tumour Growth Model

The cellular automaton model consisted of a rule-based model operating on two simulated cell populations growing on a 3D lattice. The rules and parametrisation originate from logical assumptions for tumour growth and treatment. There are four main parameters: vertical bias, cell division rate, magnitude, and length of treatment, in mathematical notation: *θ* = {*bias*, *p_divi_,λ,λ_len_*} (See Appendix 2). The model can produce six different topologies (see section 3.2.1 below).

### 2.5 Data analysis

For the calliper data set, we focus on metrics for inter-operator repeatability and the consistency of BioVolume’s linear measurements with those of callipers. For the former, we use the coefficient of variation (CV) as a measure of precision and intra-class correlation (ICC). The coefficient of variation is found as 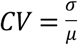, where *σ* and *μ* are the standard deviation and mean, respectively (15). For the latter, we compared the linear length and width measurements of BioVolume and callipers using a two-sample one-sided t-test with unequal variance, adjusted for multiple testing (16). For a limited number of cases, we were able to compare the calliper derived volumes to excised tumour weight. For these comparisons, we assumed the density of the tumours to be similar to that of most human soft tissues (0.90 (fat) −1.09 (skin) g/cm^3^) (17). For the evaluation of the control (*V_C_*) and treated (*V_T_*) growth curves we used Tumour Growth Inhibition (TGI):

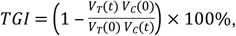

and Area-Under-the-Curve (AUC) index:

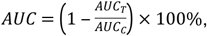

where “V” denotes volume, t time, *AUC_T_,AUC_C_* area-under-the-curve for treated and control (see Appendix 2).

A full description of the methods for the analysis of the calliper and BioVolume data can be found in Appendix 1.

## 3 Results

### 3.1 Calliper statistical review

#### 3.1.1 Inter-operator repeatability

Inter-operator repeatability is a significant challenge when measuring subcutaneous tumours (see Introduction and Graphical Abstract). It is of paramount importance that consistent and reliable measurements are made. For our evaluations we considered a Coefficient of Variation (CV) of less than 0.2 acceptable (15). Figure 2A displays the inter-operator precision for each animal model and tumour cell line. Precision was lowest for cell lines 4T-1 and A20. Across all four mouse strains for which inter-operator metrics were attainable, the distribution of precision was comparable. 59.3% of the 968 precision points scored precision values below 0.2 while 40.2% of precision values were greater than 0.2 and 0.5% lacked sufficient information to determine precision (nulls) (Figure 2B). With respect to the ICC values for volume, we obtained point estimates of 0.93 (±0.02) and 0.9 (±0.01) where measurements were made by 2 and 3 operators respectively. These values decrease to 0.64 (±0.16) when 4 operators are considered (Figure 2C).

**Figure 2:**
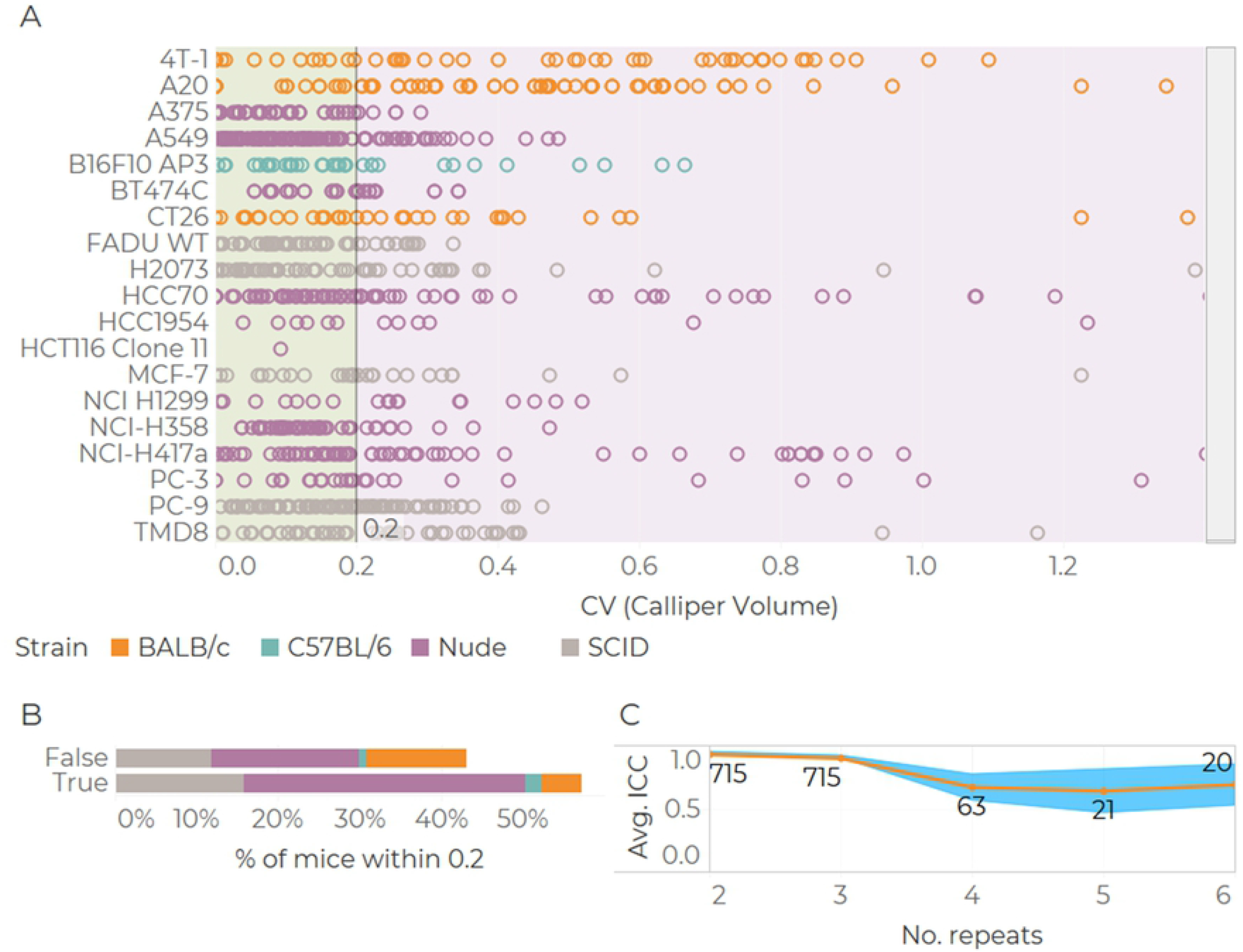
Summary of inter-operator precision and ICC in volume from callipers. Precision single values ordered by tumour model and mouse strain (A). Quantification of values within a precision limit of 0.2 (B). ICC values vs number of operators (C). Values printed on the plot indicate number of observations, dots are average ICC and shaded bars are 95% confidence intervals.

#### 3.1.2 Accuracy: volume vs weight comparison

Assuming that tumour density is *ρ_tum_* = 1*g/cm*^3^, let us define a tumour volume equivalent to *V_Eq_* = *V* · *ρ_tum_*. We find that calliper-derived volume estimates exceeded excised tumour weight in 93.7% of cases (Figure 3). The distributions of relative errors between weight and volume shows that 29.0% of tumours weigh at least half the reported volume. In contrast, only 39.8% of weights exhibited relative errors of less than 50% and only 17.4% of comparisons returned errors of less than 20%.

**Figure 3:**
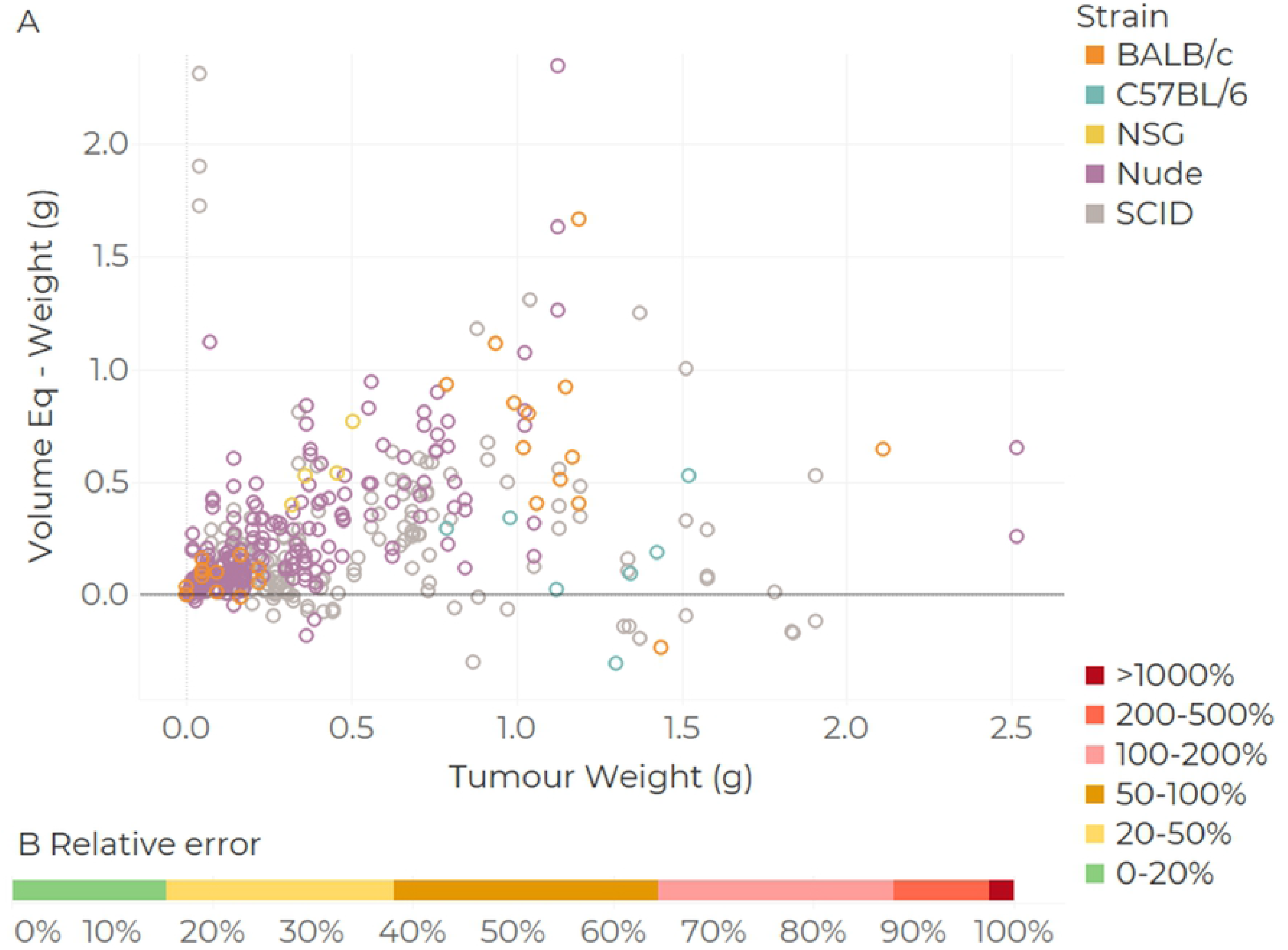
Tumour volume and weight comparison. Bland-Altmann plot (A), linear fit with 95% confidence intervals. Proportion of mice at different levels of relative errors (B, n=440). *Relative error* = (*Volume Eq. – Weight*) / *Weight*.

### 3.2 Cellular Automaton model

#### 3.2.1 Tumour morphologies

To assess the impact of using calliper measurements to estimate tumour volume we developed a Cellular Automaton (CA) model of tumour growth and its treatment (see Figure 4).

**Figure 4:**
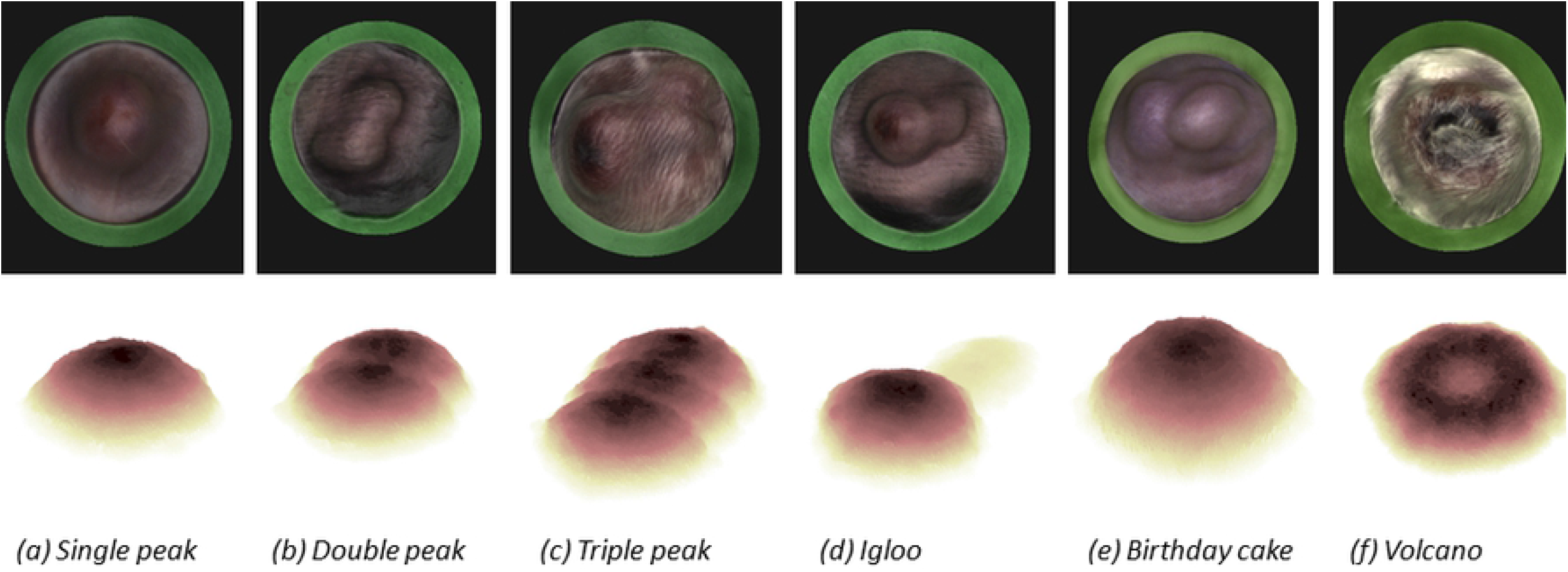
Representative examples of subcutaneous tumours exhibiting different morphologies. The top row shows reconstructions of real tumours produced using BioVolume. The bottom row shows snapshots of the corresponding in silico tumours generated with the CA model. a-c) depict tumours with one, two, and three peaks respectively. d) shows an igloo-shaped tumour. Such tumours are characterised by a main cancerous mass (typically resembling a single peak tumour) and a “tail” and can arise if the inoculating needle leaves a trail of cells when it is retracted. e) “birthday cake” tumours can be triggered by a mutation which creates a more aggressive sub-population of cells. f) volcano-shaped tumours can arise due to ulceration.

We simulated the growth of both control and treated (10 days treatment, starting on day 15) tumours for each morphology described in Figure 4. The tumours were subject to simulated calliper (SC) measurements which were then used to estimate the spheroidal volume of the tumour. These volumes were then contrasted to the actual tumour volume, which we refer to as the Ground Truth (GT). The volumes of both control and treated tumours estimated from SC measurements were significantly larger and more variable than their GT volumes across all six morphologies (Figure 5). Visual inspection of growth curves for control and treated tumours indicates that the GT volume decreased during the treatment period for all six morphologies which was not captured by the SC growth curves.

**Figure 5:**
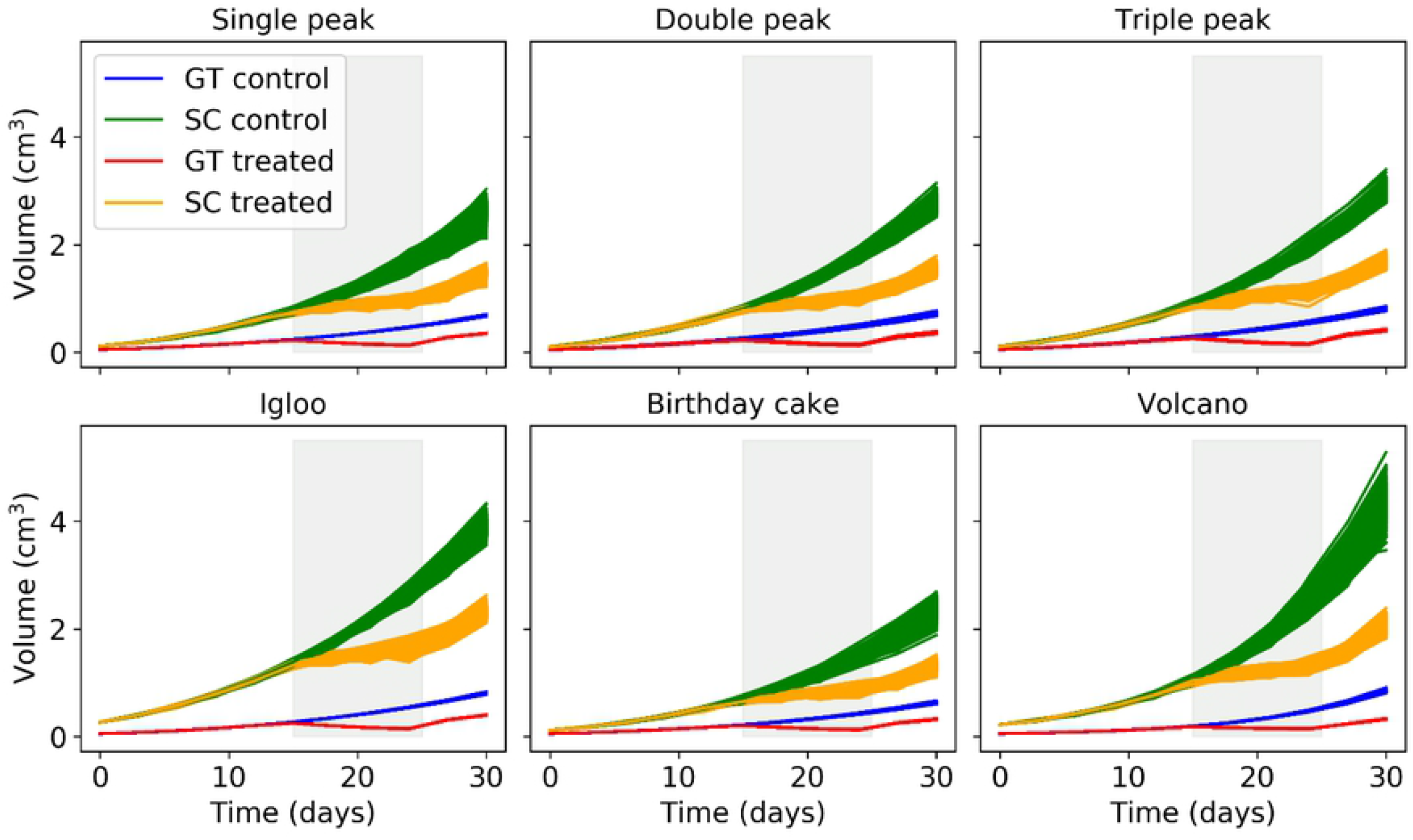
Growth curves of synthetic control and treated tumours calculated using the true volume (GT) and simulated calliper measurements (SC). For each morphology, 10^3^ growth curves were generated. The grey area corresponds to the period in which the anticancer treatment was applied; I.e., days 15 to 25.

#### 3.2.2 Treatment efficacy metrics

To determine the impact of simulated calliper measurements on the accuracy of treatment efficacy we calculated two commonly used treatment efficacy metrics using the GT and SC-derived volumes, and compared them. Specifically, we computed the Tumour Growth Inhibition (TGI) and Area Under the Curve (AUC) indices (see Appendix 2 for details). Figure 6 shows the histograms of the TGI indices computed for every pair of control and treated growth curves for every morphology at 3 different days (see Appendix 2 for histograms of combined morphologies). It is evident that the distributions are for SC have a larger standard deviation and lower level of the effect, having effectively lower statistical power. The distributions are very significantly different, overlapping only minimally. The difference is most notorious in the early treatment (see Figure 6).

**Figure 6:**
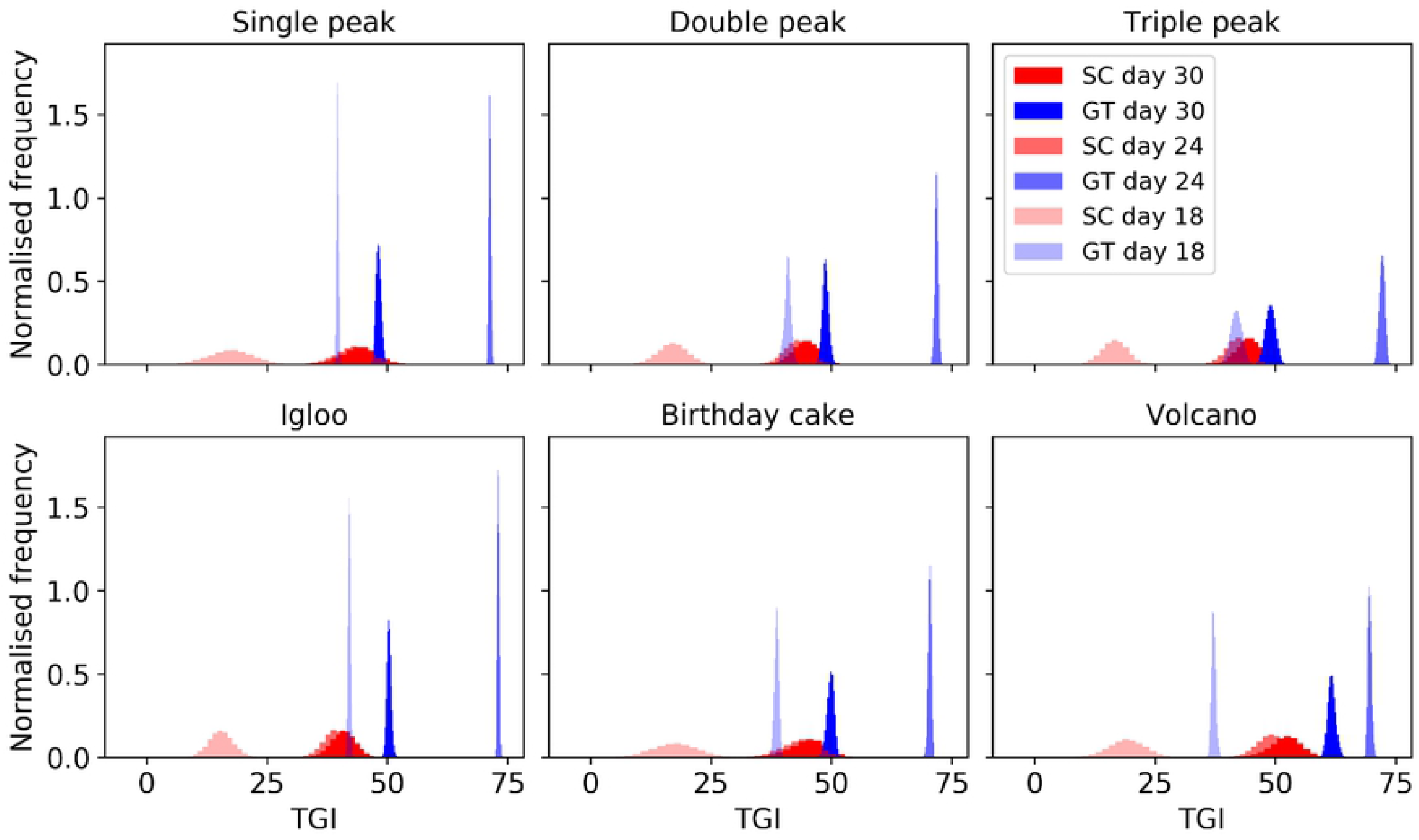
Histograms showing the Tumour Growth Inhibition (TGI) index computed for different morphologies using the true volume (shades of blue) and simulated calliper measurements (shades of red). The TGI was computed using days 18, 24 and 30 as experiment end-points.

### 3.3 Evaluation of BioVolume

#### 3.3.1 Consistency between linear measurements

We quantified the consistency between the linear length and width measurements made using BioVolume and callipers by making contemporary measurements of a given tumour using both methods and then, for each tumour, counting the number of scan measurements which fell within +/− 3mm of the calliper measurements made on the same day. These counts are displayed as histograms for both length (Figure 7A) and width (Figure 7B). Counts are organised into bins based upon the magnitude of the difference between the calliper and the scan measurements. The vertical grey band highlights values for which this difference was less than or equal to +/−3mm. The 3mm limit has been chosen as a representative figure of the standard deviation of the distributions (see Appendix 1). We found the linear scan measurements to be highly consistent with those made using callipers. For length, 88.68% of the scan measurements fell within +/−3mm of their calliper counterpart and the same was true for 90.99% of width measurements. Differences over 8mm were only observed for 2.10% of length measurements and in 0% of cases for width. In these cases, operator would outline the tumour region manually. When applying a t-test on distributions of length and width for both techniques, results are not significant with p-values of 0.330 and 0.148 respectively.

**Figure 7:**
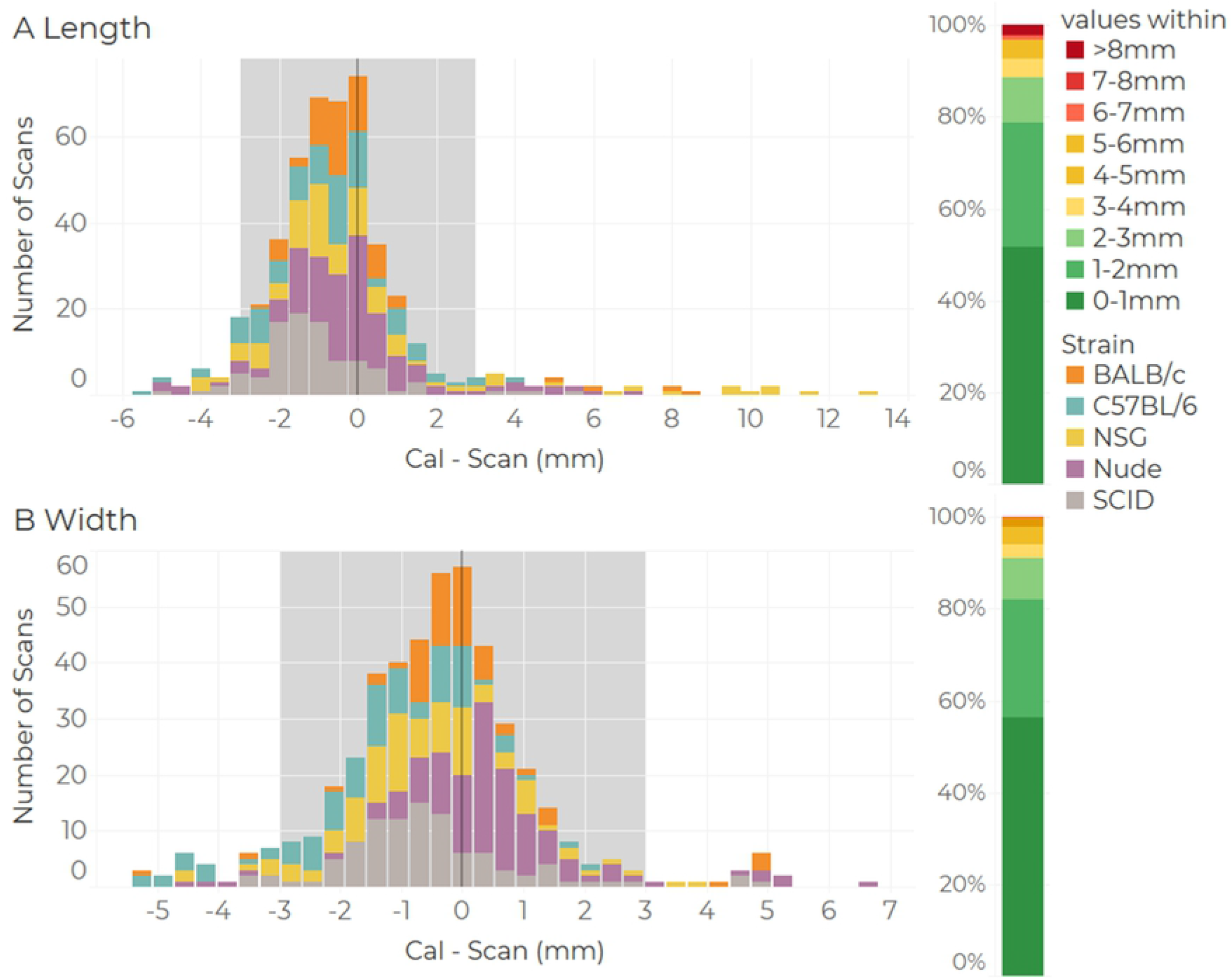
Histograms showing counts of discrepancies between calliper and scan measurements (in mm) for (a) length and (b) width of flank tumours. Counts within each bin are categorised by rodent strain. The vertical grey band highlights instance for which the difference between the scan and calliper measurement was less than or equal to 3mm. The vertical coloured bands to the right of each plot shows the number of scans falling into each range band as a percentage of the total.

#### 3.3.2 Calliper – Scan Volume comparison

Volume estimates for callipers and BioVolume were strongly correlated(*R^2^*=*0.77*) (Figure 8). Notably, BioVolume displayed a systematic tendency to estimate lower tumour volumes than callipers. This better corresponds to the lower values of observed weights relative to calliper-derived volumes observed in Figure 3. The linear trend does not exhibit any significant skewness or bias and the correlation value is high when points for which BioVolume’s linear measurements exceeded +/− 3mm of the related calliper measurements are excluded.

**Figure 8:**
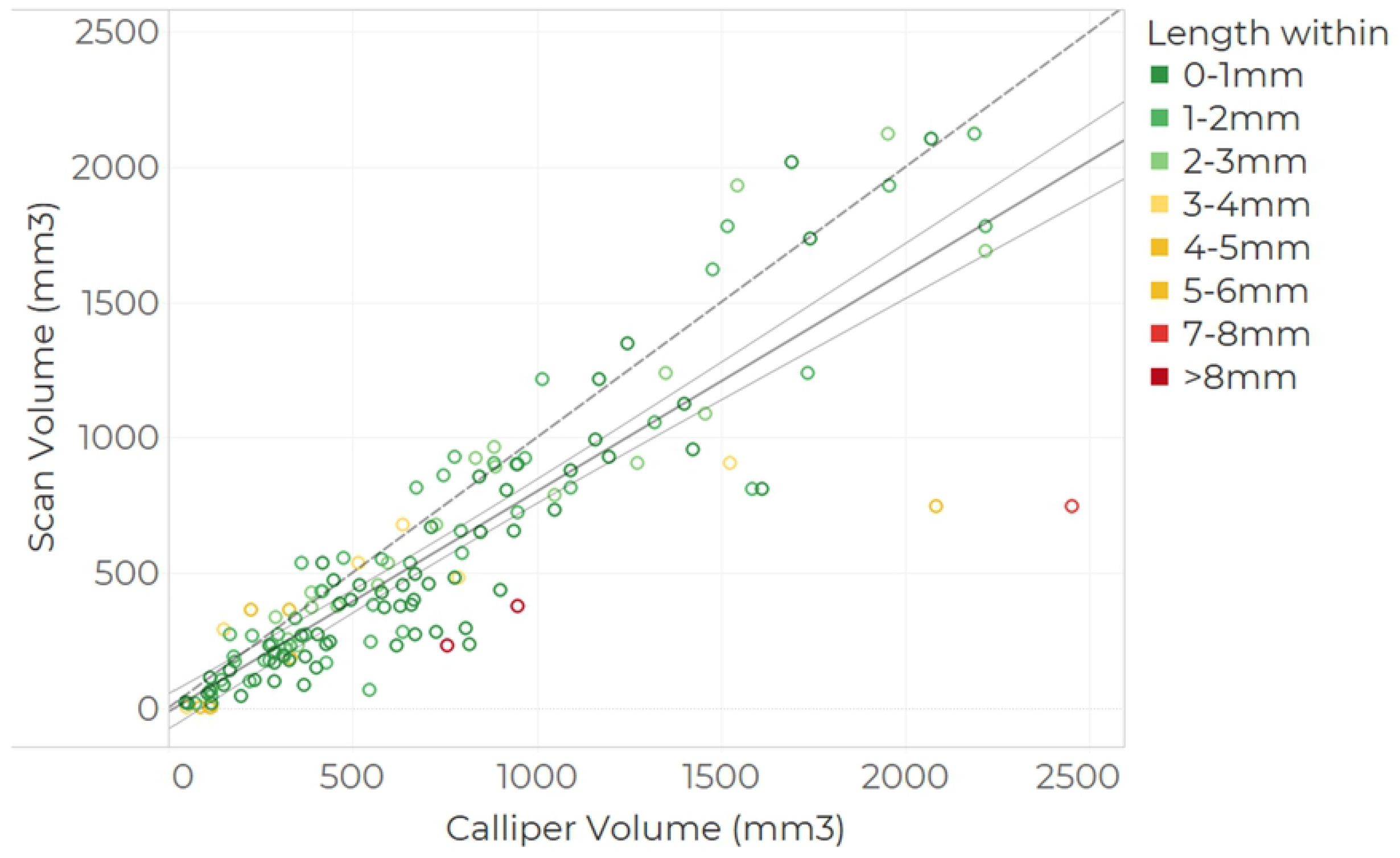
Calliper-scan volume comparison. Calliper volume was calculated using the spheroid formula, whereas scan volume corresponds to the cylindrical volume. Colour coding indicates the linear discrepancy in length between calliper and scan measurements. Linear fit is Scan Volume = (0.8146 ±0.037) · Calliper Volume + (− 14.41 ± 33.07) in mm^3^ with a correlation value of R^2^=0.77. The fit is represented by the thicker line, whereas the thinner lines represent 95% confidence interval. The dashed line shows the diagonal reference line.

#### 3.3.3 Inter-operator variability

We computed the inter-operator CV for cylindrical and spheroid volume estimates derived from BioVolume’s measurements as well as for the calliper volume estimates from dataset 1 (calliper statistical review) and 2 (BioVolume evaluation), see Figure 9. We find that BioVolume’s spheroid estimates are more precise than those of callipers, whereas the cylindrical volume, which incorporates the height of the tumour, is comparable. There are vast differences in precision between the calliper derived estimates from dataset 1 and dataset 2, with the former exhibiting greater spread and variability. Scans that were either misaligned or showed an error have been excluded mounting to 109 (38% of Dataset 2).

**Figure 9:**
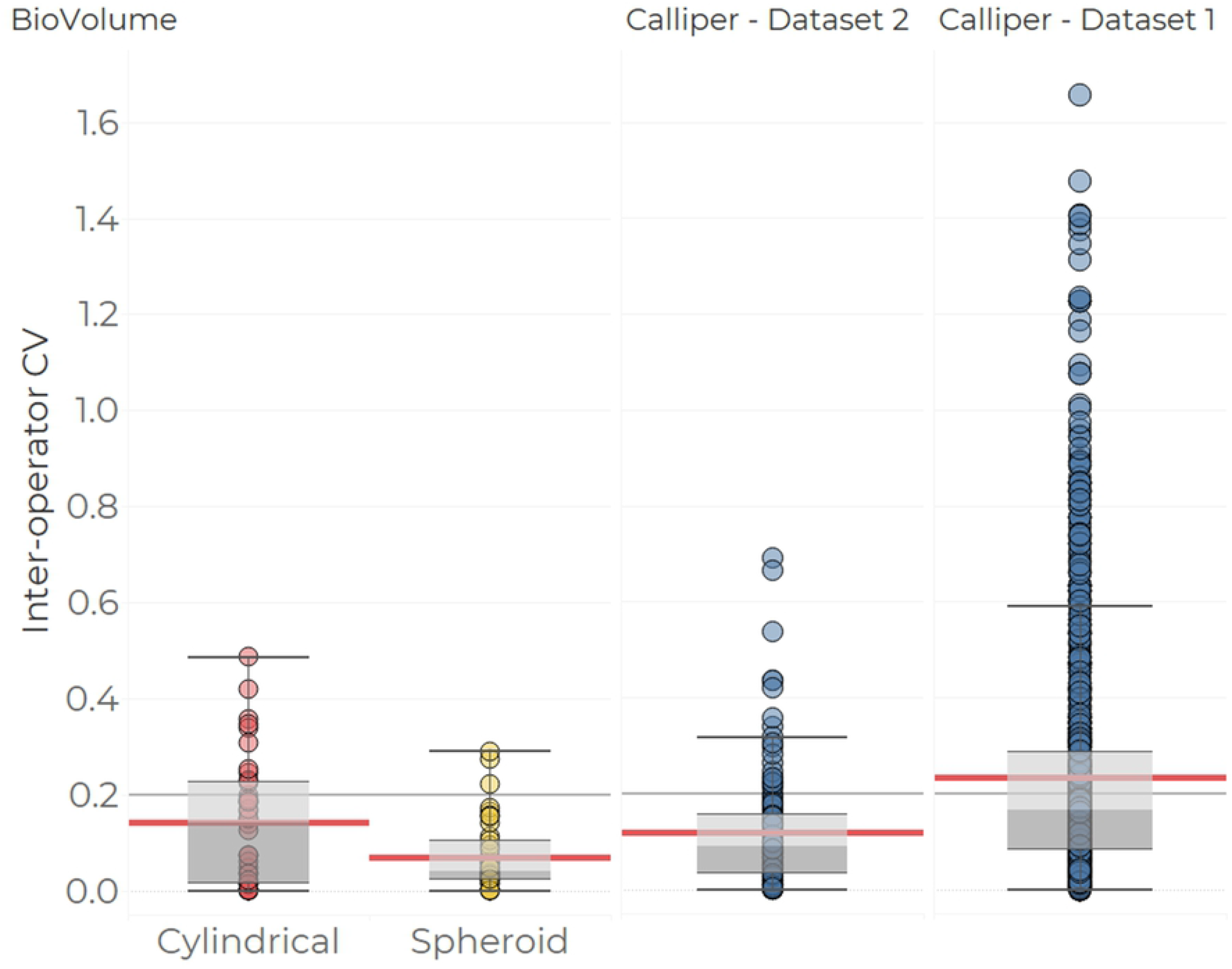
Inter-operator precision in volume estimates for callipers and BioVolume. Volume estimates for BioVolume correspond to spheroid (equivalent formula to that used for callipers) and cylindrical approximations. Calliper data is split into values from the BioVolume evaluation (left) and values from the Calliper statistical review (right, also in Figure 2). Each point captures the inter-operator CV based on two or more volume measurements made for a specific tumour on a given day. The box plots summarise the dispersal of the estimates. The main body of each box highlights the inter-quartile range while the whiskers of each boxplot encompass all values within 1.5 of the median which is indicated by the dividing line between the upper and lower hinges of each box. The light red lines reflect the mean for each category.

## 4 Discussion

In the calliper statistical review, we demonstrated that callipers are subject to high inter-operator variability, with values reaching 130% in inter-operator CV. Additionally, correlation between calliper-estimated volumes and excised tumour weight was poor (Figure 3). These metrics are alarming as tumour volume is used as a surrogate of tumour burden (weight). This may have significant implications for the evaluation of trial outcomes. Furthermore, in Figure 3 we observed that there are larger discrepancies between operators in tumour models 4T-1 and A20. This may be partly because these models are known to invade tissues locally; and partly because they were implanted under the mammary fat pad; thereby showing morphologies difficult to capture with callipers (18,19)

Our computational model clearly demonstrates that assuming tumours to be spheroid, as when making measurements with callipers, is inadequate, particularly when tumours exhibit irregular morphologies as the model outcomes were found to be morphology-dependent. First, simulated calliper measurements failed to capture the decrease in tumour volume in response to treatment (Figure 5). Second, simulated calliper measurements reduced statistical significance when comparing treatment groups as demonstrated by the TGI and AUC indices (Figure 6). Thus, when using callipers, there is an increase in the variance of measurements and a significant shift in the group mean which is sensitive to morphology (see Appendix 2). The most important model parameters were the vertical bias and the rate of cell division *p_divi_* (see the sensitivity analysis in Appendix 2). Moving forward, the model could be extended to account for different local invasion scenarios and more complex pathophysiological factors by modifying these parameters to be time-dependent. This would make the model more reflective of more complex tumour models such as syngeneic and Patient-Derived Xenografts (PDXs).

Using the prototype BioVolume scanner we were able to replicate calliper length and width measurements to within +/− 3mm in over 90% of cases (see Figure 7). Thus, both techniques produce comparable linear measurements. Volumes estimated using BioVolume were highly correlated to those estimated with callipers but were lower on average. Hence, given the trend observed in Figure 3, there is the possibility that BioVolume’s estimates will more closely correspond to excised tumour weight. This implies that BioVolume may be capable of greater accuracy than callipers, but we were unable to make a direct comparison between the volume estimates of BioVolume and tumour weight during the evaluation study. Juxtaposing Figure 3 and Figure 8, we can infer that BioVolume’s estimates, while lower than callipers, would still provide overestimates of weight. This is beneficial in practice as it minimises the risk that tumour burden limits will be exceeded. Further work needs to be performed to validate the performance of BioVolume. Specifically, direct comparisons with imaging techniques such as US, MRI or CT would provide useful insight into the accuracy of the device’s estimates.

Finally, the inter-operator variability of BioVolume outperforms that of callipers when using the spheroid formula for volume estimation. When tumour height is introduced (via the cylindrical formula, see Figure 9 and Figure S9 in Appendix 1) inter-operator variability is comparable between callipers and BioVolume. This is in part due to the fitting of the back of the mouse, the fitting of the plane to find the normal and the choice of the top point. Small variations in any of these aspects will negatively impact the consistency of height measurements. In future versions of BioVolume, we aim to improve upon these issues and to introduce a volume calculation based upon the integration of the complete surface of the tumour. We excluded 109 anomalous scans from our analysis. These anomalies arose due to system errors, misalignment of the thermal/RGB images, or motion blurring. Work is ongoing to prevent such occurrences in the future by i) improving the robustness of the code and ii) by improving the training protocol provided to experimenters using BioVolume.

BioVolume, in its current form, presents a promising alternative to callipers. It has the scope to provide accurate measurements with reduced human bias. Furthermore, measurements are fully traceable as images can be revisited at any point post-capture. Additional work is underway to improve BioVolume’s performance, and to expand its functionality. For example, machine learning can be applied to classify and characterise the stored tumour images, potentially offering additional biomarkers for treatment efficacy/toxicity and for animal welfare.

## 5 Conclusion

In conclusion, the use of linear calliper measurements for tumour volume estimation in lab animals is subject to significant accuracy and reproducibility problems which negatively affect the power of preclinical studies and animal welfare. We proposed BioVolume as an alternative to callipers which provides non-invasive, fully traceable, and more reproducible measurements with the potential to be fully morphology-independent and surpass callipers’ performance.

## 6 Acknowledgements

Thank you to AstraZeneca Bioscience Senior Management for funding this project and to all the Personal Licence holders involved in the testing. Specifically, we thank Helen Musgrove, Rebecca Whitely, Nick Moore, Emily Brough, David Simpson, Brandon Willis, Kristen Bell, Judit Espana-Agusti, Donna Goldsteen, Jane Kendrew, Graeme Smith, Chris Traher and Elizabeth Hardaker. Also, we thank the great input from Fuel3D technologies and their people to develop this technology. Special thanks to Tanya Randall, Michael Griffiths, Phalene Gowling, Fran Molina, and Chris Kane.

